# Ecological not social factors explain brain size in cephalopods

**DOI:** 10.1101/2024.05.01.592020

**Authors:** Kiran Basava, Theiss Bendixen, Alexander Leonhard, Nicole Lauren George, Zoé Vanhersecke, Joshua Omotosho, Jennifer Mather, Michael Muthukrishna

## Abstract

Social factors have been argued to be the main selection pressure for the evolution of large brains and complex behavior on the basis of data from mammals and birds. Coleoid cephalopods (octopuses, squid, and cuttlefish) have large brains, complex nervous systems and show signs of intelligent behavior comparable to that of primates, cetaceans, and birds. However, many cephalopods live largely solitary, semelparous, short lives, and many are cannibalistic, leaving little to no opportunity for parental care, complex group dynamics, or social learning. This suggests that the large brains found in cephalopods are not the result of social selection pressures. Here, motivated by the predictions of the “Asocial Brain Hypothesis”, a yet untested regime of the Cultural Brain Hypothesis formal model, we compare the relationships between brain size and social, ecological, and other factors in cephalopods. Consistent with the prediction that ecological factors should be the primary selection pressure with larger brains in more calorie-rich complex ecologies, we find that shallower and benthic (seafloor) habitats—arguably more complex environments than open-ocean (pelagic) habitats—are associated with larger brain sizes, and that measures of sociality are not. Our findings from these highly divergent evolutionary lineages, which diverged from vertebrates over 500 million years ago are not causal, but are consistent with the “Asocial Brain Hypothesis” mechanistic model that describes how ecological selection pressures can also produce large, complex brains.

## Introduction

Why do some species evolve much larger brains than others? For decades, researchers have tackled this question by identifying correlations between brain size and various ecological, social, and life-history factors. Competing hypotheses abound – from the Social Brain Hypothesis (linking brain size to social group size or other measures of sociality^1^) to ecological-intelligence^2^ and life-history explanations^3,4^. But as Shou et al.^5^ note, statistical correlations “on their own do not constitute understanding” (p. 1) We gain understanding only when we uncover mechanistic explanations for those patterns. In other words, correlations suggest possible factors in brain evolution, but by themselves they cannot tell us how or why brains evolved, nor disentangle cause from consequence among multiple confounding variables. This heavy reliance on correlational evidence has left the field with numerous plausible predictors but little in the way of unifying mechanistic theory.

Achieving deeper insight into brain evolution requires the integration of formal theoretical models that go beyond pattern-finding. Mechanistic theory can elevate our inferences by generating explicit predictions to test, thereby moving us from “just-so” correlations to causal hypotheses. Formal mathematical and computational scientific theories provide “a unifying framework”^5^ for disparate findings and can make predictions that are tested and refined through observation and serve as critical “proof-of-concept” tests of verbal hypotheses^5^. Such formal models act as rigorous examinations of the logical steps in our explanations, much like experiments test empirical hypotheses. Importantly, developing a mathematical model forces clarity about assumptions and causal pathways, often revealing whether a hypothesized mechanism could in principle produce the observed pattern. An additional advantage is that theory can tackle questions where direct experiments are infeasible: “the ability of theory to circumvent practical obstructions of experimental tractability… is a benefit that should not be underestimated”^6^ (p. 5). Even when we cannot manipulate evolutionary pressures in real time, we can use models to explore how evolution might have proceeded under different conditions. In sum, formal theory provides the mechanistic and predictive backbone needed to interpret correlational data in a more meaningful way. That is, we can evaluate the degree to which different models can produce empirically discovered correlations between features of species within a taxa.

Within the study of brain evolution, new theoretical frameworks are emerging to unify the diverse drivers of brain expansion. The motivation for the present study is one such theory developed by Muthukrishna, et al.^7^ - the Cultural Brain Hypothesis (CBH). The CBH proposes that brains evolve principally to acquire, store, and manage information. Rather than focusing only on social group size or any single factor, the CBH posits that larger brains are selected for their capacity to handle more information – whether gained through individual (asocial) learning or social learning. This theory captures how cognitive demands (learning skills, innovation, memory) co-evolve with life history and sociality, providing a unifying framework for brain-size variation across taxa. Crucially, the CBH model yields distinct scenarios or “regimes” of brain evolution. The theory was developed to understand the cultural and social pressures typical of humans and primates, hence the term “Cultural Brain”. In the first paper proposing the theory, the CBH predictions were shown to reproduce the correlations between primate brains, group size, innovation, social learning, mating structures, and the length of the juvenile period. The model was subsequently tested with whales and dolphins using a new database created for the purpose of testing the model^8^. Given the nature of marine research, the data was less rich than with primates, but here too, the model was consistent with brain size, social structures, and cultural behaviors across cetaceans.

The formal model from which the CBH was derived unexpectedly also predicted a regime with a pathway to big brains driven not by social learning but by asocial information acquisition in complex environments. We call this the “Asocial Brain Hypothesis” (ABH). In this paper we use the same protocol for creating a database and testing the CBH predictions with cetaceans to test the ABH predictions with cephalopods.

The ABH extends the logic of the CBH to species with minimal social learning: it argues that even in largely solitary animals, intense ecological challenges and individual learning opportunities can favor increased brain size as long as learned skills allow an animal to acquire sufficient calories to support the larger brain. In other words, the calorie richness and complexity of the environment rather than social complexity becomes the primary selection pressure on brains in asocial lineages. The details of the model are discussed in the next section. In the present study, we put these theoretical ideas to the test by examining brain evolution in coleoid cephalopods – a group of animals that offer a compelling asocial contrast to the usual social vertebrate models.

Octopuses, squids, and cuttlefish have some of the largest brains (relative to body size) among invertebrates and exhibit complex behaviors on par with those of birds and mammals. Most cephalopods present a paradox for social brain theories: they are generally asocial creatures. Many octopus and squid species are short-lived and semelparous (dying after a single reproductive event), largely solitary, and even cannibalistic, which leaves little opportunity for extended parental care, social transmission, social tolerance, intricate dominance hierarchies, or cultural learning. Their large brains clearly did not evolve for managing stable social groups. Why, then, did cephalopods evolve such big brains? The Asocial Brain Hypothesis was not developed with cephalopods in mind and therefore does not fully capture some aspects of this group, such as the reproductive cycle. Nonetheless, as Servidio et al.^6^ explain for the role of abstract biological models, it serves as a proof of concept of mechanisms that could generate observed correlations and therefore a guiding framework for data collection and analysis. The ABH makes explicit how if social factors are minimal, then ecological factors, particularly availability of calories with trial and error learned knowhow (for how to navigate the complexity to unlock those potential calories) should emerge as the main correlate of brain size.

Motivated by this insight, we compiled comparative data on brain size and a suite of ecological, behavioral, and social variables across 79 cephalopod species. We find that the cephalopod data strongly support this prediction. Species inhabiting calorie-rich complex environments have significantly larger brains, whereas measures of sociality show no such relationship. These results align neatly with the idea that ecological complexity, not social complexity, has been the primary driver of cephalopod brain expansion. It is important to note that we interpret these correlations as consistent with a theory that offers a mechanistic explanation, but not as direct causal claims based on the data itself. That is, the positive correlation between benthic living and brain size does not causally demonstrate that a complex habitat selects for bigger brains in real time – many unmeasured factors could be at play and other models could in theory produce the same predictions. However, such a finding is both motivated by and consistent with the ABH’s mechanistic causal scenario in which calorie richness and ecological challenges create selection pressures favoring cognitive investment. In this way, the correlations serve as empirical support for theoretical predictions in the absence of experimental manipulation, even if there may be other verbal models (see next section) and formal theoretical models yet to be developed that show the same pattern of results.

In addition to our overall findings, our study illustrates how formal models and comparative data can work hand-in-hand to advance theories of brain evolution. By showing how even an asocial lineage can evolve large brains under the right ecological conditions, we expand the explanatory scope of brain-evolution theory beyond primates and other social animals. More broadly, this work underscores the value of combining mechanistic models with comparative analyses. Such an approach helps unify disparate findings and encourages the development of general frameworks applicable across life’s diversity. In moving beyond simple correlations toward theory-driven tests, we take a step closer to understanding how and why brains evolve along different paths.

### Prior research and alternative hypotheses

The evolution of large brains across the animal kingdom has been a subject of extensive research, with various social, behavioral, developmental, and ecological factors being proposed as key drivers of this energetically costly adaptation^9–12^. Comparative studies have identified a range of predictors for larger brains, including group size, social structure, sexual competition, parental care, length of the juvenile period, predation pressures, diet, locomotion, and sensory capabilities^13–17^. Historically, such research has been biased towards groups similar to humans, both taxonomically (e.g. primates and other mammals) and behaviorally, with high levels of sociality (e.g birds). Therefore, social factors have become the dominant explanatory predictor, after initial work by Jolly^18^ and Humphrey^19^ argued for social complexity as a driver for intelligence among primates, and the hypothesis gained support on the basis of data from mammals and birds^16,20–22^. The social brain hypothesis (SBH) in particular, was developed to explain the evolution of large brains in primates^1^, and argues that larger brains evolved to cope with the complex demands of social group living, such as cooperation, deception, and competition^23–25^, as well as pair bonding and social cohesion^21,26^.

The SBH has been tested among various taxa with nuanced results. Some findings have implicated the quality of social relationships rather than group size. Among birds, for example, species with small social groups (5-30 individuals) seem to have larger relative brain sizes than solitary species or those forming large groups^26–28^. An analysis of sexual competition and brain size in bats found larger brains among species with monogamous rather than promiscuous females^29^. A study on African cichlid fishes included sex-specific analyses and found that females providing sole parental care had larger brains than those in species with biparental care, with no effect of care type on male brain size^30^. A comparative study of cetaceans found a quadratic relationship with group size, where the largest-brained species (and those with larger social repertoires) formed mid-sized social groups rather than smaller/loose aggregations or megapods^8^.

In other taxa, no relationship or even a reverse relationship has been found between social measures and brain size. Among carnivorans, an analysis using fossil data indicated that, contrary to a previous analysis of mammalians^20^, sociality was not linked with increased encephalization across clades and this relationship was likely driven solely by canids^31^. A study of a rodent group, African mole-rats, showed larger absolute brain size and neuron numbers among solitary species, with the authors highlighting the entanglement of body size with both brain size and the evolution of group living^32^. Among non-avian reptiles, relative brain size was not correlated with habitat complexity, and in contrast with findings in primates, solitary species had larger brains than social ones^33^.

Most of the research effort has been on vertebrates. Social insects are a notable exception with some studies exploring the relationship between sociality and brain size. Due to the nature of labor division in many of these species, sociality could be expected to have an inverse effect on brain size; however, differences in expected cognitive load due to variation in how specialization takes place in a colony– e.g. by age where an individual progresses through all tasks in some groups, changes in how labor is divided as a colony grows in size, and variation in colony size over time and within species– complicate this prediction^34,35^. This very different expression of sociality, as well as the presence of high intraspecific variation in relative brain volume and the contribution of particular cellular mechanisms to behavioral variation apart from volumetric measures of brain size, may explain the inconsistent results for tests of the social brain hypothesis among insect groups^34,35^.

Taken together, these disparate findings highlight how expressions of social behavior with differing cognitive demands - e.g. dominance hierarchies in primates, cooperative breeding in birds, and colonial societies of related individuals in insects - across and within different taxa complicate any clear relationship with behavioral complexity, cognition, and brain size. A recent meta-analysis of the Social Intelligence Hypothesis (here encompassing the Machiavellian Intelligence Hypothesis and the SBH) found broad support for a “significant positive relationship between sociality and cognition across interspecific, intraspecific and developmental studies”^36^. The authors note that despite criticisms of measures of brain size and cognitive ability, there was not a significant difference in effect sizes across studies with different measures of cognition, or between studies of primates vs. non-primates. However, they also note that the vast majority of studies have been carried out on primates, other mammals, and birds, and that insects - the only invertebrates in the studies surveyed - were highly underrepresented given the number of taxa. Therefore, it remains unclear the extent to which sociality is linked with cognition outside these heavily-studied taxonomic groups.

Two other potential explanatory variables for the evolution of large brains are ecology, and life history and development. Research on the importance of ecological complexity in driving brain size has focused on the cognitive abilities required for innovation, problem-solving and foraging for difficult-to-extract or high-energy foods and the larger brains that can be supported by ecologies with more calories and energy availability. An early study by Clutton-Brock and Harvey^37^ found differences in brain size (relative to body size) across primates according to ecological differences, including diet (e.g. folivores and frugivores) and home range size. A later study of primates tested measures of ecological vs. social variables^38–40^, finding frugivorous diets (requiring spatial memory and extractive foraging and providing higher energy) were a better predictor of larger brains than social group size, while there was no support for the SBH. A comparative study of mammals and birds^38–40^ found that while experimental measures of self-control were predicted by absolute brain size, they were not strongly linked with residual brain size or at all with social group size, casting doubt both on the link between sociality and cognition and the validity of residual brain size as a measure of cognition. Another study found that brain size among primates was predicted by manipulation complexity, which was also linked with tool use and extractive foraging^38–40^.

The ‘cognitive buffer hypothesis’ focuses on life history and development, arguing that the behavioral flexibility afforded by large brains will allow organisms to respond to socioecological adversity, with the fitness cost of additional developmental time offset by reduced extrinsic mortality, resulting in overall longer lives and time for reproduction^41^. There has been some support for this hypothesis, with studies of mammals^41–43^, birds^41–43^, and amphibians^43^ showing a coevolutionary relationship between large brains and longer lifespans. Kaplan and Robson^4^ (2002) use a formal model and empirical data to demonstrate a coevolutionary relationship between brain size and longevity in primates, a correlation also shown in parrots^44^. However, a recent analysis of cetaceans showed a negative relationship between brain size and lifespan^45^.

This body of research has been largely empirical, yielding disparate and sometimes contradictory relationships between a variety of variables^13,20,42^. One way to unify this disparate body of work is through the use of general theories that formally describe the complex co-evolutionary relationships between all relevant variables.

### An attempt at unifying models of brain evolution

General theories that formally mathematically or computationally model the causal pathways and coevolutionary relationships can shed light and make clear predictions. One such model was developed by Muthukrishna et al.^7^. The analytic and computational models (see Figures 1 and 2 below) were developed to incorporate sociality, ecology, and life history into a single framework. It builds on previous work, including the verbal theories of the Cultural Intelligence Hypothesis^46,47^, but formalizes assumptions about the relationships between brain size, organismal fitness, and energy requirements. The model revealed two social regimes and an unexpected third asocial regime for the evolution of large brains. The first set and second set of predictions were for social and highly social animals. The “Cultural Brain Hypothesis”, described the social learning path to larger brains, which has been tested with primates^3^ and cetaceans^8^. The “Cumulative Cultural Brain Hypothesis” revealed a narrow set of parameters within the social learning path that would lead to an autocatalytic takeoff consistent with the human pathway characterized by an ever-increasing reliance on culture and technology (see also^48^). The predictions in the third asocial regime, which we label the “Asocial Brain Hypothesis” (ABH), were untested and therefore motivated the present research on asocial species with a large literature and data collection. Before we present the results, we summarize the assumptions and predictions of the model.

**Figure 1.**
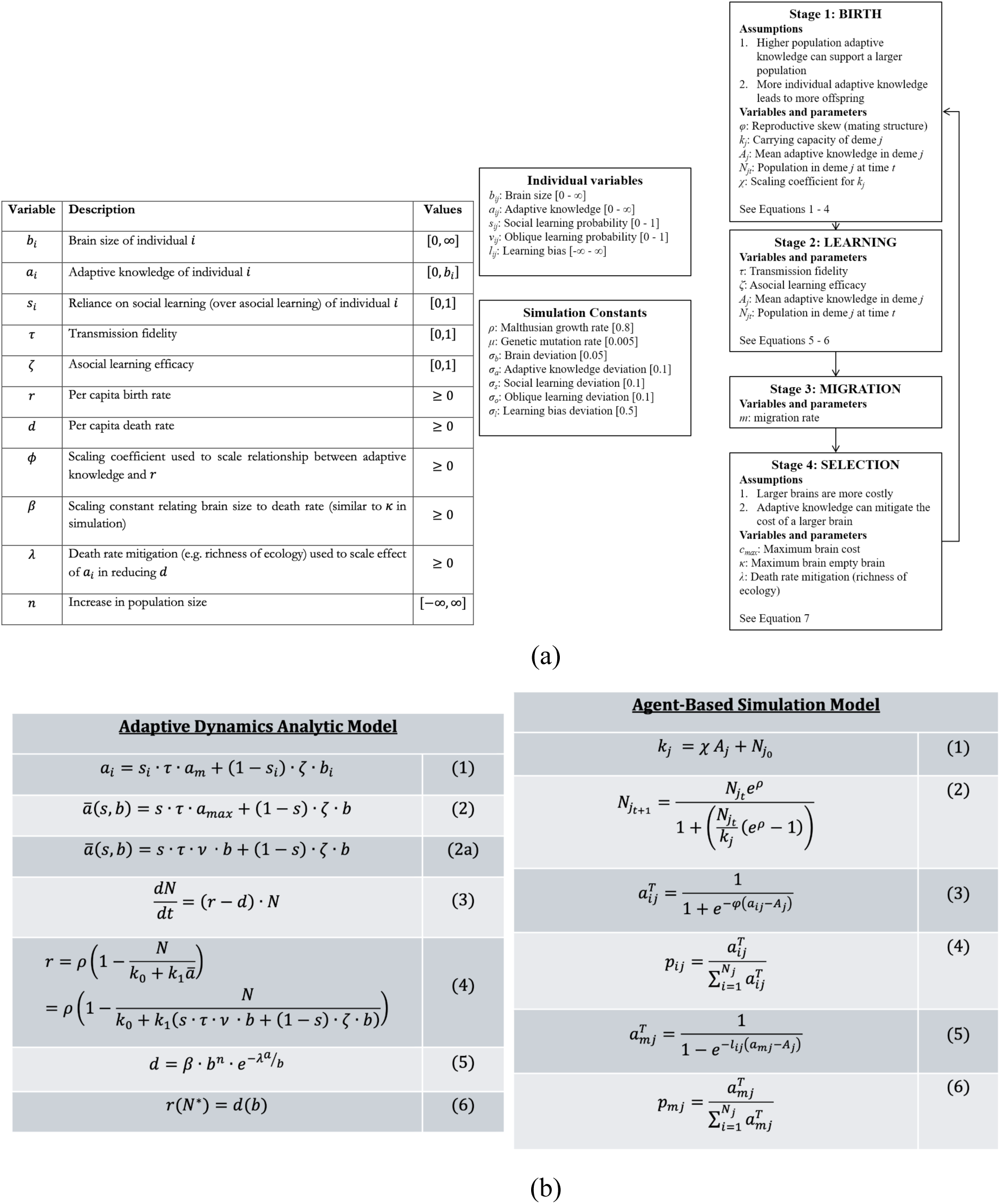
(a) Variable definitions for the analytic model, with individuals *i* represented by brain size *b_i_*, adaptive knowledge *a_i_*, and reliance on social over asocial learning *S_i_* (b) Variable definitions for the agent-based simulation model. The mathematical relationships between variables for the analytic model (left) and agent-based simulation model (right).

**Figure 2.**
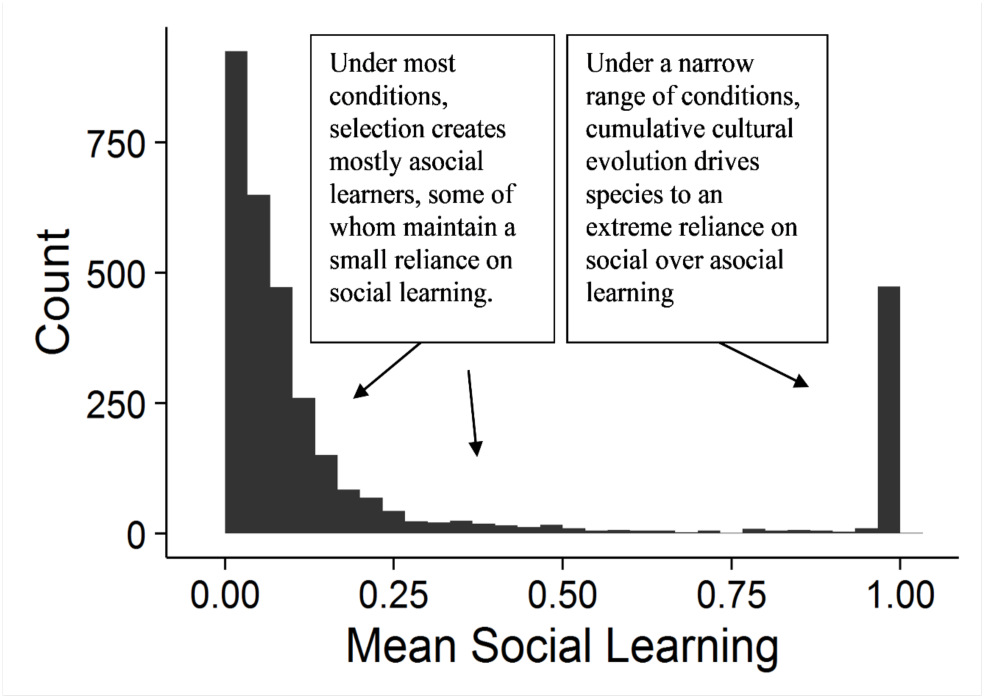
The different equilibrium regimes for reliance on social learning that emerged in the simulation. Most conditions select for individuals primarily reliant on asocial learning (left side), while a small range of conditions may create individuals primarily reliant on social learning (right side). A narrow set of conditions lead to an extreme reliance on social learning (far right). These mechanisms identified by the model that lead to these three regimes are the Asocial Brain Hypothesis, Cultural Brain Hypothesis, and Cumulative Cultural Brain Hypothesis, respectively.

The formal model (from which the Cultural Brain and Asocial Brain Hypotheses are derived) had three key assumptions from which all other relationships emerge:

1. Larger and more complex brains are more costly than less complex brains because they require more calories, take longer to develop, and have organizational challenges. Therefore, increasing brain size/complexity decreases an organism’s fitness; an organism should have the smallest brain possible that allows it to thrive.^1^ BUT
2. A larger brain correlates with an increased capacity and/or complexity that allows for the storage and management of more adaptive knowledge.^2^ AND
3. More adaptive knowledge increases an organism’s fitness either by increasing its number of offspring compared to conspecifics and/or by reducing its probability of dying before reproduction. Adaptive knowledge can be acquired asocially, through experience, trial and error, and causal reasoning, or socially, by learning from others.

The predictions derived from these assumptions were modeled using two approaches. An analytic approach using an adaptive dynamics evolutionary model^49^ to show how brain size, adaptive knowledge, and reliance on social learning varies as a function of transmission fidelity in social learning (which may be a function of social tolerance, theory of mind, etc), asocial learning efficacy (which may be a function of learning opportunities, physical ability to manipulate the environment, etc), and survival returns on adaptive knowledge (via calorie availability as say a function of sunlight and water).

The model leads to 3 key predictions, only the third of which is relevant to the present study on asocial animals:

(1) Increased reliance on social learning requires high transmission fidelity (relative to the ability to generate knowledge by oneself).
(2) Extreme reliance on social learning requires access to a range of models with different amounts of adaptive knowledge.
(3) A greater return on adaptive knowledge (affected by calorie richness of the environment) increases brain size.

The complexities of co-evolutionary dynamics and more complex social learning strategies were difficult to derive analytically, so instead, a simulation was used to capture these complexities. Figure 1a below shows the variables for the analytic and agent-based simulation models, and the stages of the simulation model. Figure 1b below shows the mathematical relationships between the variables in the analytic and simulation models. The analytic and simulation solutions can be found in Muthukrishna et al.^7^.

The simulation reproduced the results of the analytic model with nuancing around social learning strategies. A sweep across the range of parameters suggested that most species have limited social learning (top left of Figure 2), but that within this regime, there was still a pathway to larger brains. This “Asocial Brain Hypothesis” (ABH) suggests that such “trial and error” learning animals would ignore conspecifics and therefore have a weaker, non-existent, or even negative relationship between brain size and measures of sociality (such as group size or intensity of social interaction) and would have to reduce the length of a juvenile period spent exploring rather than exploiting knowledge. The logic behind this is that trial and error learning is less efficient than social learning because the space of possible behaviors and interactions to be learned is much larger than simply learning from a filtered set of socially accumulated actions discovered by other group members; for example, it’s faster and easier to be taught something than figure it out on your own. For animals relying on asocial learning, the pressure to begin exploiting learned knowledge is greater and as a result, we expect that the evolution of larger brains in asocial animals to be primarily constrained by the complexity and available calories in their ecology – there is no corpus of social knowledge that incentivizes a longer learning period.

### Overlap with previous theories

Although the ABH formalized how asocial learning behavior and life history could interact in certain ecologies to produce large brains, it is not the first to suggest this relationship. Previous research has likewise emphasized the role of ecology in behavioral and brain evolution. Among these, Wyles et al.^50^ advanced the “behavioral drive” hypothesis, where animals in a population capable of socially transmitting innovative behaviors can exploit new niches, leading to novel selection pressures that further favor adaptive behaviors and subsequent physiological evolution. However, their focus was on terrestrial vertebrates and discussed social transmission, while the ABH focuses explicitly on asocial animals that learn independently through trial and error. Kaplan and Robson^4^ focus on long-lived, social groups– specifically primates– and find a coevolutionary relationship between brain size and lifespan. They employ a formal model in combination with empirical data and emphasize the importance of ecological conditions in shaping the relative energetic costs of larger brains.

### Cephalopods as a valuable comparative case in brain evolution

With a last common ancestor with humans and other vertebrates over 500 million years ago, cephalopods not only have flexible bodies, but also flexible cognition and behavior. For instance, many species of cephalopods exhibit a variety of foraging and predator-avoidance strategies^51^. These strategies, such as camouflage, navigation, mimicry, and tool use can be flexibly adjusted to nearby species or other contextual cues^52,53^. Although they do not display the cohesion and complex sociality of many mammalian and bird species, communicative body patterns are used widely across the squid and cuttlefish in mating contexts, where aggressive visual signaling as well as tactical deception have been observed in a range of species^54–56^. Many species, especially the well-studied *Octopus vulgaris*, display cognitive abilities such as tool use, navigation and problem solving as well as many types of learning^57^. Playful behavior, an otherwise rare occurrence in the animal kingdom, has also been observed in some octopuses^58^.

Cephalopods, therefore, represent a well-studied, asocial, independent evolutionary origin of intelligence from that of the vertebrate animals we often view as possessing advanced cognition, such as primates, cetaceans, and birds. While gregarious behavior is observed in squid, cephalopods tend to live largely solitary, semelparous, short lives^51,59–65^. Many are cannibalistic, a barrier to social tolerance let alone sociality^66^, including complex group dynamics or social learning.. In our current sample of 79 species where brain data is available, at least incidental cannibalism was observed for 21 species with data largely unavailable for the remainder. A previous review of a broader set of species found cannibalism in 34 species, associated with various factors including population density, body size, stress, and prey availability^66^. Cephalopods also exhibit little to no parental care or pair-bonding, often dying after first reproduction^67–69^. As these characteristics run counter to established theory and empirical findings from other, more social, vertebrate taxa, cephalopods are a promising animal group on which to evaluate prominent hypotheses on the possible evolutionary drivers of brain size^70–72^.

As discussed above, explanations focusing on the coevolution of sociality and intelligence do not translate well to cephalopods, particularly the notoriously solitary octopuses. Researchers have instead focused on explanations involving predation pressures to these soft-bodied organisms and their reliance on extractive foraging in complex oceanic environments^64,65,73^. Packard^74^ extensively documented convergences in cephalopod form and behavior with fish, highlighting the common environment and resource competition that has led to similarities, including in brain development and cognition. Cephalopods would have faced selection pressures to compete for resources, avoid predation by, and predate on fish. Research on cephalopod cognition has illustrated their reliance on exploration and problem-solving, seeking out and integrating information from their environments for extractive foraging, capturing prey, and interacting with conspecifics^51^. In particular, the extensive repertoire of complex defensive behaviors possessed by extant cephalopods highlights the likely significance of predation in their cognitive evolution^75^. Phylogenetic studies incorporating data from the fossil record support theories that competition with marine vertebrates during the Mesozoic spurred the evolution of an internal shell and greater locomotion and speed^76^, which allowed them to colonize a much greater range of ecological niches as well as exposing them to greater predation pressures, increasing selective pressures for sophisticated camouflaging and cognitive flexibility^64^.

Cephalopods also contrast with cetaceans (the marine group previously tested for the cultural brain hypothesis^8^), through occupying a far broader range of ocean habitats ranging from shallow coastal water, the open ocean, and the sea floor at a wide range of depths^77^. Overall, shallow benthic (seafloor) habitats can be considered more calorie-rich, energy available, and biodiverse environments than pelagic (open water) habitats. However, these measures depend on factors such as geographic scale, latitude, depth, substrate, and specific organismal lifestyles (e.g. daily vertical migration from the depths to the surface)^78,79^. Additionally, much remains to be discovered about organisms inhabiting the deep sea, open ocean, and other environments^80,81^. To capture the variety of ecological selection pressures these habitats impose, we include several environmental variables including benthic vs pelagic habitats, depth in meters, and latitude (described further in results and methods).

To test the ABH, we systematically surveyed the cephalopod literature to identify and capture a wide variety of qualitative and quantitative data on extant coleoid cephalopod species’ brain and body size, physiology, ecology, life-history, sociality and behavior. We read all 3933 papers in the Zoological Record and extracted all available data on 115 variables (see Supplemental Material for full list) for all 79 species for which comparable adult brain size data are available from the year 1866 to July 2020. We also added papers published up to 2024 which contained new brain data. This data revealed large gaps in what we do and do not know about extant coleoid cephalopods. Here we report the results of the 3 variables for which we had full data: ecology (as measured by benthic vs. pelagic habitat), depth (minimum and maximum, with minimum more reliable as maximum depth is more difficult to confirm), and sociality (as measured by categorizing species as solitary or gregarious, with an additional category of tolerant/aggregative in sensitivity checks). The availability of these data allowed us to compare ecology to sociality. The results of other analyses, including those relevant to the theoretical predictions, are less reliable. All other variables with sample sizes are reported in the Supplementary Information (SI).

## Results

The Asocial Brain Hypothesis predictions as they apply to cephalopods were pre-registered (https://osf.io/nbwsx). Here we present the key results with the most complete data, comparing ecology against sociality, with the full set of results in the SI. All analyses are Bayesian multilevel models with total central nervous system (CNS) volume as our measure for brain size as the outcome variable. Missing covariates were imputed during model fitting, meaning that available information was used to estimate unobserved values. Where multiple data points were available for the same species, we ran it with the maximum brain size value in the main analysis and with all brain size values in sensitivity checks (reported in the SI) - these did not affect the results. We also ran other robustness and sensitivity checks, including considering the maximum, mean, and minimum of some variables such as depth and age at sexual maturity to assess consistency of observed effects across models (SI). To account for the scaling of brain with body size, each model also includes as a predictor dorsal mantle length (ML) as a standard measure of cephalopod body size^82^. All continuous variables are logged and standardized (mean 0, standard deviation of 1) and all models incorporate phylogenetic relationships as a correlation matrix to account for evolutionary relatedness between species; for further details on model specification and variable operationalization, see the Methods section. All analysis code and data are available in the supplementary. All analysis code is also openly available at https://github.com/kcbasava/ceph-brain-evolution and data are also available at cephdata.com and will be posted on Figshare.

For each model, we report the point estimate for the mean of the posterior distribution and the lower and upper bounds of the 95% Credibility Interval (CI). The posterior distribution contains the probability of parameter values conditional on the model and data. The result plots report marginal posterior predictions with covariates held at their mean. A headline summary of the results for the main variables is presented in Figure 3 and Table 1. As mantle length is a reliably large and positive predictor of brain size (a posterior mean point estimate of ∼0.75 across models on a logged and standardized scale) with a known biological cause, for ease of interpretation, we interpret the coefficients of our predictors benchmarked relative to this estimate as this is plausibly the largest effect size we could observe (i.e., body size should be the strongest predictor of brain size).

**Figure 3.**
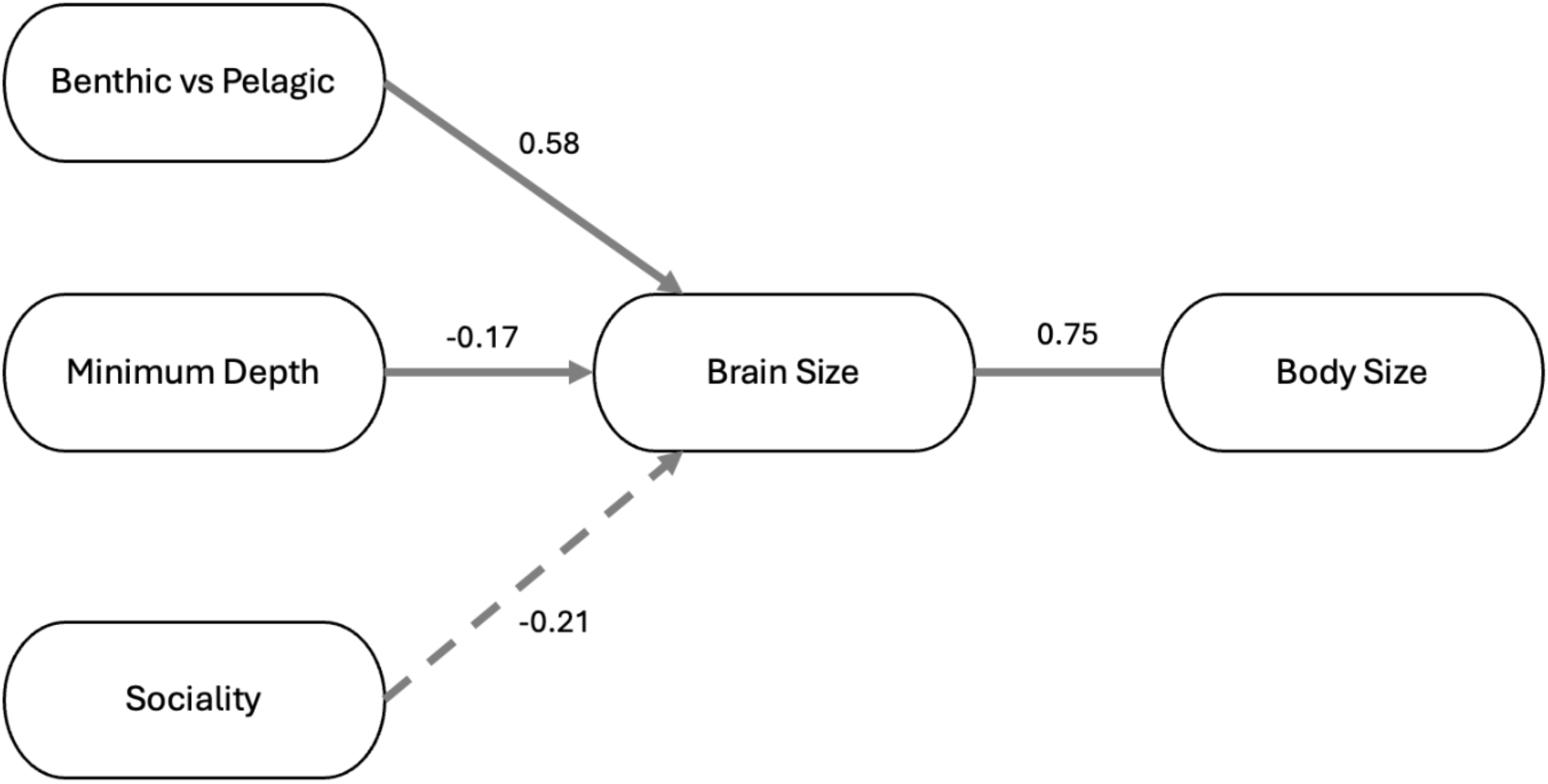
Strongest predictors of brain size found in our analysis. The dashed line from sociality indicates that the credibility interval included 0. Note that these are the main effects with phylogenetic and research effort controls.

**Table 1.** Summary of the results for our main predictor variables, indicating the strength of the relationship relative to that of body to brain size, the direction of the relationship and consistency/lack thereof with the ABH, and our subjective assessment of the reliability of this relationship. Remaining results using variables with more missing data is reported in the SI.

| Variable | Strength of relationship relative to ML:brain relationship | Sign of relationship | Results consistent with ABH | Strength of evidence* | Number of species with data on focal predictor |
| --- | --- | --- | --- | --- | --- |
| **Benthic (vs. pelagic) habitat** | $\sim 2/3$ | positive | yes | strong | 79 |
| **Depth** | $1/4$ (minimum depth)<br>$1/8$ (maximum depth) | negative | yes | medium | 79 |
| **Sociality** | $\sim 1/3$ | negative | yes | medium | 79 |

\*Informal strength of evidence assessment based on amount of missing data, source data quality, effect size, CI ranges, and accuracy of variable definitions, as discussed in the main text.

The relationship between brain size and habitat was two thirds as large as the relationship between brain size and body size; the relationship with minimum depth was a quarter, maximum depth was an eighth, and the relationship with sociality was a third, but in the negative direction.

### 1. Ecological complexity

For asocial learning species such as cephalopods, the ABH predicts a positive relationship between brain size and richer ecologies across all regimes^7^. This prediction emerges in the model because richer ecologies with more calories can support larger brains for both social and asocial learners. Additionally, this relationship is expected to be reinforced by a greater behavioral repertoire leading to more access to calories consistent with ecological explanations for brain evolution.

In our statistical models, ecological complexity was operationalized through: habitat type (benthic, pelagic); minimum and maximum depth in meters and broad categories (shallow, deep, ontogenetic or daily migration); latitude; dietary breadth, measured through the number of taxonomic categories consumed (e.g. crustaceans, gastropods); and number of predators, also measured through the number of taxonomic categories (e.g. other cephalopods, bony fishes).

#### 1a. Habitat type

We ran an analysis considering habitat as a binary variable (benthic vs. pelagic). The effect of benthic habitats on CNS was positive (0.58 [0.08, 1.07]). To test the robustness of this effect - as lifestyles are not always easily divisible into benthic and pelagic - we also considered a three-category measure of habitat (pelagic, benthic, and variable/near bottom/demersal, which under the binary measure is classified as benthic). The results still revealed a positive effect for near bottom/demersal habitat species (0.64 [0.09, 1.20]) and for non-bottom benthic species (0.54 [-0.02, 1.09]). The estimates for pelagic were somewhat negative in both analyses (-0.42/-0.41 [-1.60/-1.56, 0.79/0.77]). We plot these relationships in Figure 4 below.

**Figure 4.**
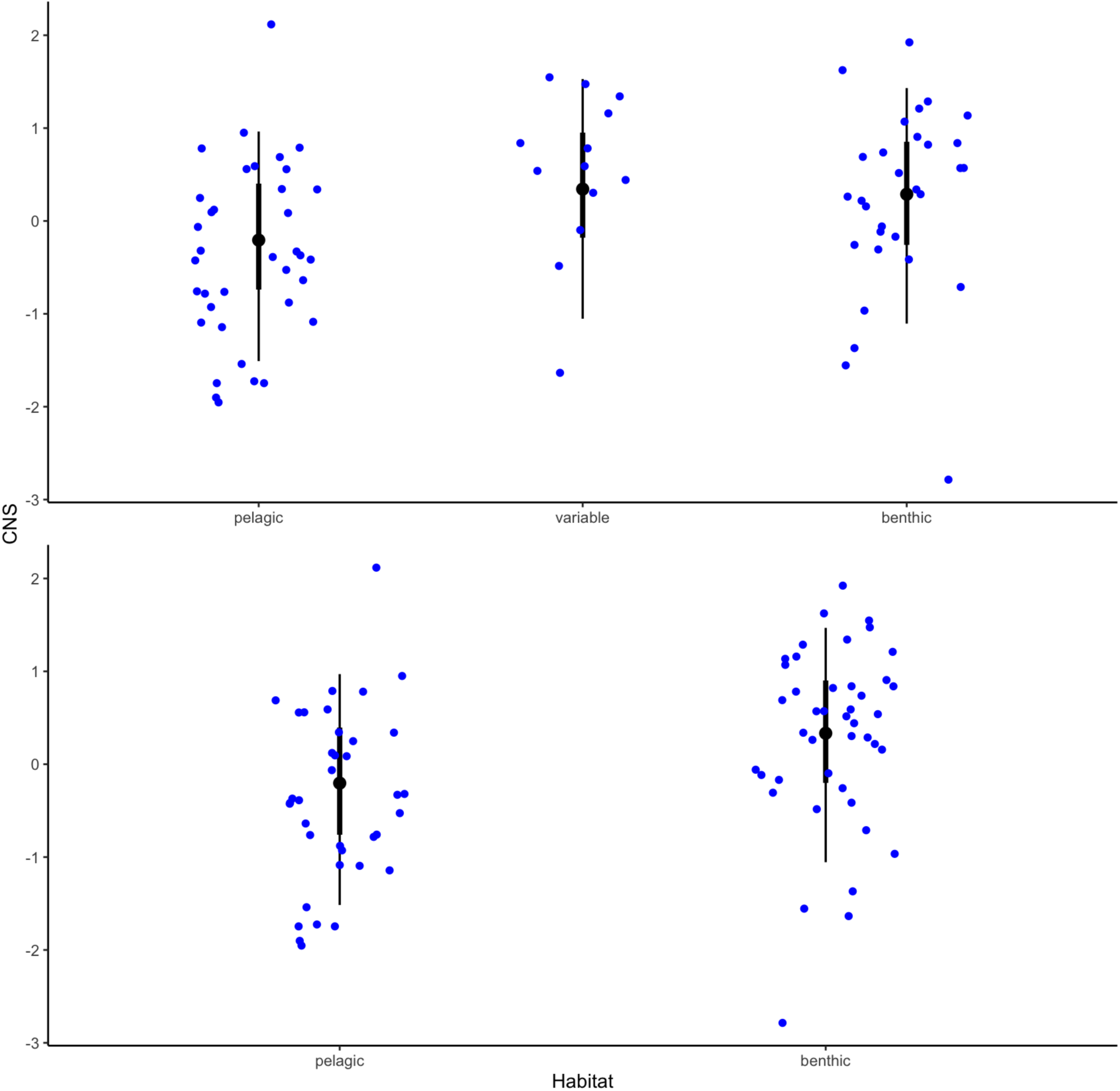
Plot of results from analyses predicting brain size from habitat coded into three or two categorical types. Bars show predicted CNS (medians with highest density continuous intervals) and blue dots observed data for three-category habitat (top) and binary habitat categories (bottom).

While correlational, the results of the analyses conducted with the first version of habitat codes are consistent with the predictions of the ABH with selection for larger brains in more calorie-rich, complex habitats. However, this is complicated by some species displaying elements of a pelagic lifestyle in addition to interacting with the benthos. Depth range as a potential proxy for ecological complexity did not positively predict larger brains in our analyses (it did show a negative effect, but this was likely driven by higher maximum depth in species with the largest depth ranges). The distribution of depth ranges in meters for each species in the dataset is shown in Figure S2, colored by habitat and ordered in ascending order of relative brain size (CNS/ML). In particular, the plot illustrates how species with relatively larger brains tend to live in benthic or variable habitats.

The phylogenetic analysis of Lindgren et al.^77^ indicated convergent morphological changes in cephalopods with shifts to different oceanic habitats. These included the presence of photophores for bioluminescence in pelagic cephalopods, accessory nidamental glands to protect eggs for species which attach them to substrate, and corneas among benthic taxa for protection from sediment. It is logical that selective pressures of different habitats would likewise influence brain evolution across cephalopod taxa.

#### 1b. Depth

Depth measures in our dataset are somewhat imprecise given that many species have wide and/or variable ranges or engage in vertical migration, and due to the potential biases in measurement methods (e.g. different capture methods in fisheries studies). However, we attempted a rigorous analysis using both minimum and maximum recorded depths, as well as a categorical measure that took into account life history differences in depths occupied. Noting that minimum measures are more reliable as it is easier to measure how shallow a species occurs than how deep, minimum recorded depth had a negative effect (-0.17 [-0.28, -0.04]). Maximum recorded depth also had a negative effect (-0.10 [-0.26, 0.04]), though the CI included 0.

Considering depth as a 4-factor categorical variable, categorizing species into shallow water, daily vertical migration, ontogenetic migration (usually descending as they grow from paralarvae/juveniles to adults), and deep sea also indicated lower CNS for ontogenetically migrating species (-0.85 [-1.35, -0.24]) and deep-sea species (-0.70 [-1.21, -0.20]). The estimate for shallow-water species was centered around 0 (0.15 [-1.00, 1.31]) and for daily vertical migration slightly negative (-0.26 [-1.82, 0.82]). These estimates are plotted in Figure 5 below.

**Figure 5.**
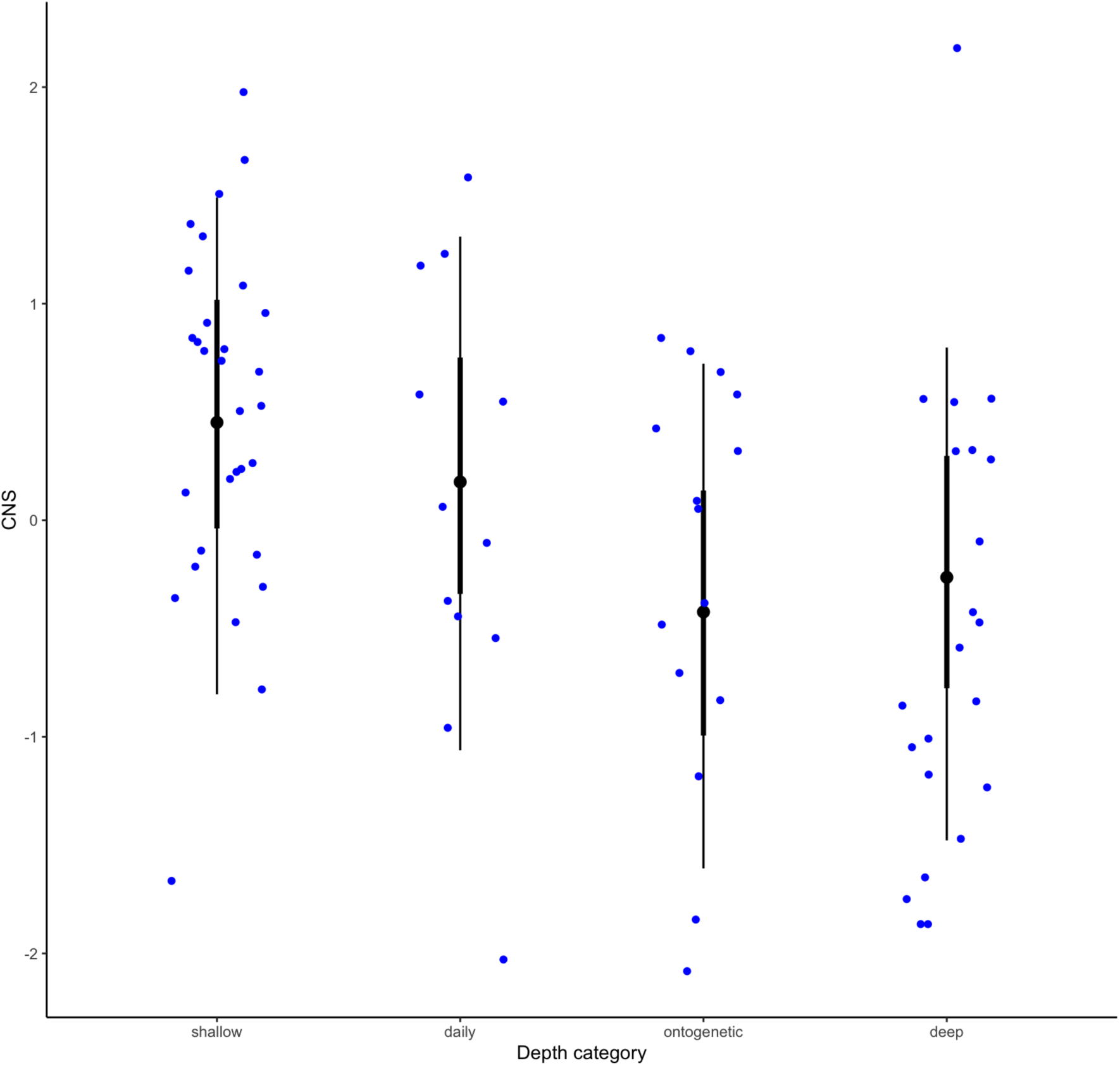
Plot of results from analyses predicting brain size from depth measured in four categories. Bars show predicted CNS (medians with highest density continuous intervals) for different depth categories and blue points observed data. From left to right: shallow water, daily migration, ontogenetic migration, deep water. CNS is logged and standardized.

In summary, the effect of depth is negative, consistent with our predictions, assuming that shallow habitats tend to be more complex than deep-sea ones due to the presence of sunlight and greater energy availability.

### 2. Sociality

The ABH predicts a weak, non-existing, or even negative relationship between brain size and sociality for cephalopods, as asocial learners do not benefit from a larger group in the way social learners do (larger groups have more models to learn from). This prediction is one of the key differences between the SBH and ABH. Consistent with this prediction, whether species were solitary or gregarious was not positively associated with brain size (Figure S3). Sociality instead had a somewhat negative effect (-0.26 [-0.60, 0.17]), although the credibility interval crosses 0, within the range of the ABH predictions. We also ran an analysis using three categories for sociality rather than the binary measure, with an additional ‘in-between’ code meant to capture species that appear tolerant of or would associate with other individuals close by (coded as solitary in the binary variable). These include some species that have been observed aggregating in the wild or displaying pair bonding behaviors such as *Abdopus* species^85,86^ or that appear unusually tolerant of proximity in lab conditions such as *Rossia macrosoma*^87^ and *Sepia bandensis*^88^. The results were broadly similar for social species strictly coded (-0.21 [-0.58, 0.21]). Tolerant species had a slightly higher estimate for brain size (0.37 [-0.02, 0.79]), but interpretation of this result is complicated by the small number of species in this category (11 total) and the heterogeneity of characteristics that could classify species as falling between solitary and gregarious. These results are shown in Figure 6 below.

**Figure 6.**
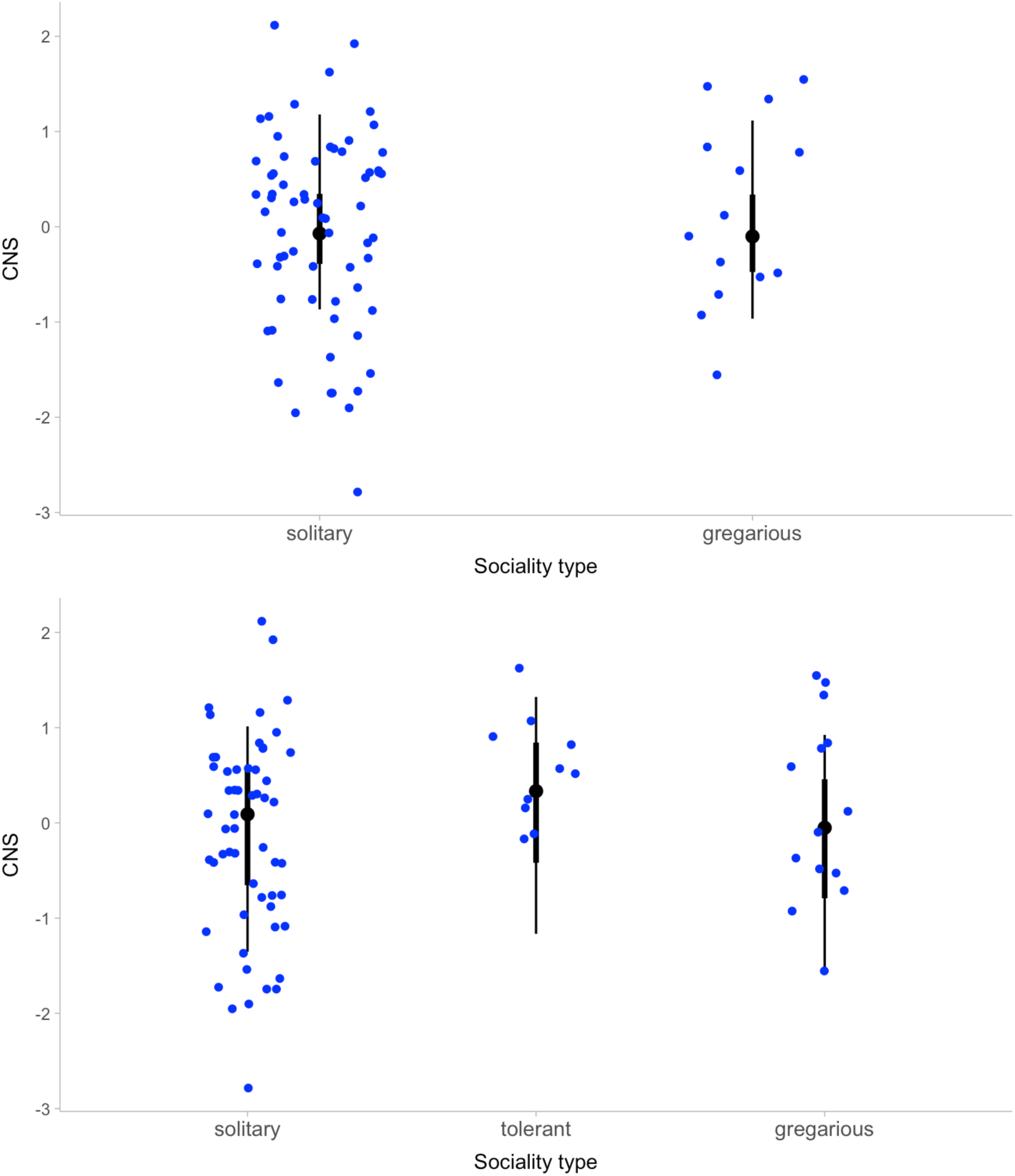
Plot of results from analyses predicting brain size from sociality measured as a binary (top) and 3 categories (bottom). Bars show predicted CNS (medians with highest density continuous intervals) for different sociality categories and blue points observed data. CNS is logged and standardized.

**Figure 7.**
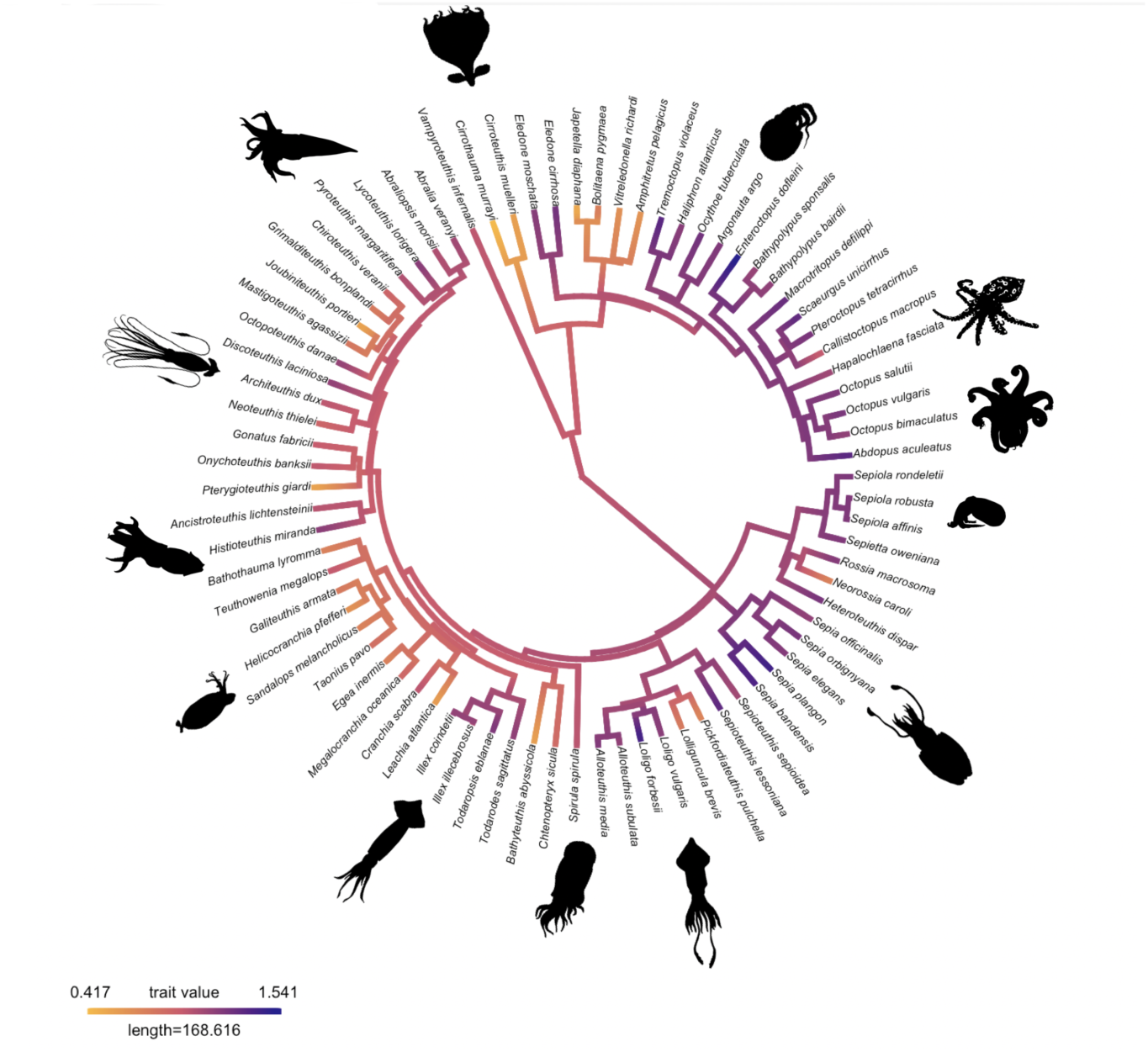
Ancestral state reconstruction of relative brain size (CNS/ML) on a log scale. Images from phylopic.org (Keesey 2024).

A supplementary analysis was also run to look at sociality only among the 54 decapodiforme species (squid, bobtail squid, and cuttlefish) in our dataset. The effect remained negative (-0.24 [-0.76, 0.25]), indicating that this effect was not driven only by larger brains among octopuses, which are predominantly solitary. A corresponding analysis among octopodiformes was not possible, as none were coded as gregarious.

Our sociality findings contrast with many of the predictions and findings from mammals and other social vertebrates. Sociality in cephalopods is strongly phylogenetically conserved, with schooling present exclusively (though not universally) among the decapodiformes. Solitary and social species can generally be clearly distinguished; however, there are many aspects of sociality such as group sizes and schooling dynamics in the latter that have not been adequately observed or described in the literature. Cannibalism, common among at least some cephalopods, is another factor that may have obstructed the evolution of richer social lifestyles through inhibiting the possibility of any social trust between conspecifics^66^. Additional field research is needed to draw more fine-grained conclusions about how sociality interacts with cognition and brain size in cephalopods.

## Discussion

We present the largest and most comprehensive database of cephalopod species where brain data is available. Analyses of this dataset show phylogenetically controlled relationships that are consistent with ecological theories of brain evolution, but not consistent with social theories^1,89^. Measures of ecological complexity, but not solitary vs. gregarious lifestyles, predicted larger brain sizes in our dataset. This was the case both across species in all three major groups and within squids and cuttlefish, which do display some levels of social grouping and interspecific interactions. Habitat in particular emerged as the most reliable predictor of brain size in cephalopods, with shallower-living and benthic species tending to have larger brains. Since shallower and benthic habitats are more calorie-rich and biodiverse relative to deep sea and pelagic habitats. For instance, pelagic food webs tend to have lower species diversity than do benthic ones, possibly leading to greater specialization among benthic foragers^80,90^. This finding is consistent with the idea that environmental variability selects for greater cognitive capacities and larger brains^91^. Although we did not evaluate different lobe sizes, our findings are also consistent with Ponte et al.’s^92^ assessment of cerebrotypes in relation to phylogeny, development, and environmental niches which emphasized the importance of habitat and life strategies in shaping brain variation across species. Overall, these findings are consistent with hypotheses that have emphasized how large brains may coevolve with the ability for innovative and complex behaviors, allowing animals to survive in variable environments and cope with ecological challenges^77,91^.

### Limitations and future directions

The main limitation of this analysis is the sparsity of cephalopod data available in the literature, including for species for which brain data is available. Our dataset includes comparative quantitative brain data for 79 species. These are all the species for which brain data is available, but amounts to roughly 10% of the total number (around 820) of extant cephalopod species. To maximize the use of the data we do have, we used a Bayesian multilevel modeling approach so that no data was discarded. Nevertheless, the amount of data does affect the size of the Credibility Intervals which both emphasizes the strength of the ecological findings, but urges caution for the more limited data availability driving the analyses in the SI.

For example, our measure of predator breadth, which is an important aspect of ecology, suffers from missing data, overgeneralization of taxonomic categories, and difficulty with accurate measurements for many species. More data would be required to create a higher resolution measure of calories and complexity, incorporating temperature, latitude, and oxygen levels affecting diversification, competition, and predator-prey interactions^83,84^ (see SI for further discussion). However, the categorical measure of deep-sea lifestyles and ontogenetic migration to the depths both showed lower estimates of brain size. Overall, further research on specific environmental aspects of different depths, as well as how variables like predation pressures and prey availability differ for cephalopod species inhabiting benthic or pelagic habitats^75^ will likely shed more light on ecological selection pressures on brain size.

This contrasts with the complete data in habitat type, which is far easier to measure and classify. A far greater amount of lab and field data would be needed to properly test the predictive value of variables like behavioral innovation rate, which has been shown to predict brain size in birds and primates^14,93^. While our results similarly support some aspects of the cognitive buffer hypothesis^94^, i.e. selective pressures for cognitive ability from a variable environment, this predicts that large brains will preferentially be selected in longer-lived species^41^, which does not apply well to short-lived coleoid cephalopods. Further research will help clarify the aspects of these different environments that serve as selection pressures, and how these interact with variables such as predation pressure and diet that were less conclusive in our analyses.

Another possible limitation is that by constraining our analyses to species with presently available quantitative brain data, we risk a selection bias if these particular ‘brain data’ species are exceptional in relevant characteristics compared to non-sampled species. However, the present dataset spans a wide variety of species that are diverse in size and habitat and covers a wide range of families across the major lineages (squid, octopuses, and cuttlefish) of the coleoid cephalopod phylogeny. It therefore seems unlikely that brain size directly influences the probability of sampling, which would be the worst case scenario from a selection bias perspective.

A further limitation of the current study was the single measurement of brain and body size for the majority of species, which means that intra-species variability was not taken into account and idiosyncrasies of the individuals that happened to be measured might have affected the validity of our results. To mitigate this, we ran sensitivity checks by including the additional brain and mantle estimates for the 36 species in our dataset with additional (one or two) brain and mantle length measures. The results of these analyses are summarized in the SI and are consistent with the estimates from our main analyses. There is also the possibility that as methods used to capture brain size differ between older and newer analyses, leading to inconsistencies between data we combined. As with ecological and behavioral data, future research producing additional measurements for brain size would be valuable to further evaluate processes of brain evolution.

Finally, as our analyses used whole brain size as the focal outcome, they did not take into account measures of brain structure, organization, and neuron numbers which have been suggested to relate to variation in behavioral repertoires and complexity in cephalopods and other species^95,96^. Among cephalopods, this includes research on relative lobe size and how these link with ecological factors^69,97,98^. Ideally, additional work measuring these factors among different cephalopod species will make an analysis incorporating these other potentially important measures possible in the future.

In summary, although these results are correlational, they are consistent with ABH model predictions. Modeling is essential to theory development for the study of complex systems^99^, and can help make sense of the variety of empirically discovered relationships reported in the literature. Through our analyses, we found evidence for the importance of ecology, rather than sociality, for brain evolution in cephalopods. Our findings contrast with the relationship between sociality and brain size among primates and other vertebrate species, drawing attention to a previously neglected group in this literature and illustrating the potential significance of ecological complexity as a selective pressure. We also collated a large comparative dataset of ecological, behavioral, and physiological traits from an exhaustive survey of the literature on cephalopods that can be expanded upon and used for future comparative research. We hope that this dataset and approach motivate more specific formal theories that capture the evolutionary history and phenotypic constraints of cephalopod physiology and life history as building blocks of a fully general theory and theoretical framework of brain evolution^100^.

## Methods

Prior to the development of the cephalopod database we present here, the most comprehensive comparative study on cephalopod brain size to date was published as part of a 2007 PhD dissertation by Borelli^101^ and in a subsequent journal article by Ponte and colleagues^92^. This ambitious and useful contribution had limitations with regard to broader research efforts on comparative brain evolution:

1. The dataset expresses brain size in relative terms (specifically, the size of each brain lobe is computed as a proportion to the whole central nervous system) and does not include absolute brain and body size measurements, which makes these data unsuitable for comparative regression analyses.
2. The dataset does not include social, behavioral, or life-history variables necessary for directly evaluating predictions of the ABH.
3. Due to the nature of available cephalopod phylogenies, Ponte et al.^92^ were forced to discard data on more than half of their 78 species when running phylogenetically-controlled analyses.
4. Upon reviewing Borelli’s primary brain data sources^69,97,98^ and their subsequent use in the literature, we found inconsistencies that ultimately required us to conduct a careful, systematic re-evaluation of all data (refer to Supplementary Information for details).
5. The dataset contains brain measures from the juvenile specimens for a few species. As cephalopod brains are still maturing during the juvenile period^102,103^ and possibly during adulthood^104^, it is not appropriate to use these for comparative analyses with measures from adult specimens.
6. Considerable advances in empirical work on cephalopods, including additional brain data measurements, have been made since Borelli’s pioneering 2007 PhD data collection.

The present study overcomes these limitations. The dataset we present in this paper was developed with three main goals^73^: (1) to assess key hypotheses on the evolutionary drivers of brain size in cephalopods, starting with the Asocial Brain Hypothesis (ABH); (2) more generally, to move toward a theoretical reconciliation of empirical findings from comparative brain size studies in other animal groups; (3) provide a dataset to the cephalopod research community for further analyses and theory-building. Drawing on the predictions of the ABH as well as previous brain evolution literature, here we empirically assess whether measures of ecology, age at sexual maturity (i.e., length of juvenile period), sociality, and behavioral complexity predict brain size in coleoid cephalopods.

### Data collection

Brain and body size data were taken mainly from Wirz^98^ and Maddock and Young^97^, updated by Nixon & Young^69^, combined with a few newer species-specific brain data estimates^105–107^, including Chung et al.^105,106^ and Montague et al.^107^. We used brain size measurements from adult specimens of identifiable species (79 total). Data on behavior, ecology, physiology, and other traits were collated from a systematic review of articles written on each species on Web of Science, taking into account alternate species names. The bulk of the data was captured up until November 2020, which was followed up by a more targeted data collection effort for particular species and variables up to 2024. We also captured and coded data from major books on cephalopod ecology, physiology, and behavior^67,69,108–111^.

Due to a lack of data on pre-adult stages, we coded our variables focusing on adult brain and behavior. Also due to limitations of the data, we did not distinguish between observations of male or female individuals except in cases of sex-specific variables (e.g. male courtship displays).

### Variable operationalization

Brain size was operationalized as the total central nervous system (CNS) volume including optic lobes (in millimeters^3^) from the sources listed above. Body size was operationalized as the mantle length from the same specimens in millimeters. In the analyses reported here, we use the maximum recorded total CNS volume and, when available, its associated mantle length size. As sensitivity checks, we re-ran main models with available supplementary CNS data (see https://github.com/kcbasava/ceph-brain-evolution/tree/main/sensitivity-checks).

We operationalized the length of the juvenile period/age of sexual maturity through the oldest age at which a species is not observed to be sexually mature measured in number of days, with lifespan (also in days) included in the models for which this was a predictor. We analyze both the maximum and minimum recorded ages. Sociality was measured as a categorical variable with species coded as solitary (avoiding other conspecifics except for mating), tolerant/aggregative (an in-between category with individuals sometimes observed as congregating outside of reproductive contexts and appearing tolerant of nearby individuals), or gregarious (usual or obligate schooling). This was coded as a binary variable in the main analyses with ‘tolerant’ species coded as solitary/0. Sensitivity checks were run with the three-category measure.

Species were characterized as having benthic or pelagic habitats, with benthic defined as animals living on and/or closely interacting with the seafloor and pelagic animals primarily living in the water column. After further review of these variables, it was decided to create a third category for species observed interacting both with the seafloor and swimming in the water column (demersal/in-between) that would otherwise be classified as benthic. Depth was measured through the minimum and maximum depths in meters at which adult individuals of each species had been recorded, as well as a 4-category variable which classified species as living in shallow waters, undergoing daily vertical migration, migrating to the depth as adults (i.e. ontogenetic migration), and living in the deep sea.

Behavioral complexity was measured through (a) combined social behaviors and cognitive measures, (b) number of defense/antipredator behaviors, and (c) number of foraging behaviors, each used as predictor variables (full description in SI). This measure was intended to capture behavioral complexity both as measured through lab studies of abilities such as learning, memory, and problem solving, as well as through field observations of interactions with conspecifics in the wild, e.g. with deception in mating contexts and communication abilities, for species which have not been studied in one or the other context. Unfortunately, this variable was too sparse to yield results we felt were robust, hence they are not included in the main text and we do not draw theoretical conclusions from them. As we did preregister and run analyses with behavioral complexity as a predictor, for transparency they are included in the SI.

Researcher effort was accounted for by including variables for (1) number of articles published on Web of Science for the species name (or alternative names) and (2) articles read during coding of the dataset. The former was used as the standard variable for research effort while the latter was used for variables more likely to be biased due to frequency of observation (i.e. count variables including dietary breadth, number of predators, or foraging strategies).

### Statistical models

All analyses are Bayesian phylogenetically controlled, multilevel linear regression models and were run using the *brms* package in *R* ^112^, an interface to the probabilistic programming language *Stan* ^113^. They generally follow the operationalization of Bayesian multilevel regression models from McElreath^114^ as implemented in brms by Kurz^115^. The multilevel structure of the model accounts for the non-independence of the data from species in our dataset through incorporating information on phylogenetic relationships. All models included brain size (as total CNS volume) modeled as a function of body size (as mantle length). Other variables were included as derived from adjustment sets from a directed acyclic graph (DAG) depending on the focal predictor. DAGS describe a hypothesized causal model through nodes (variables) and directed edges (relationships)^116^. The code describing the DAG, including all variables and hypothesized causal paths, are available in the *R* scripts at https://github.com/kcbasava/ceph-brain-evolution. All variables except for habitat, categorical measure of depth, and sociality were standardized (mean centered with a standard deviation of 1) and log-transformed prior to analyses. Priors were chosen to be weakly regularizing to help prevent the model from overfitting to the data^114^. A consensus phylogeny created from the results of the phylogenetic software (discussed in detail in the Supplementary Information) was incorporated as a covariance matrix following the implementation in Kurz^117^. Models were fitted according to the following structure:

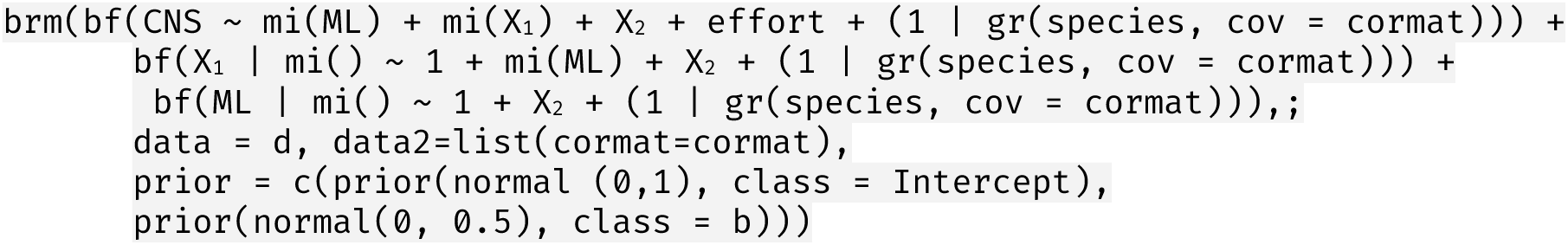

where CNS is total brain volume, ML is mantle length , effort is research effort, X_1_ is a predictor or predictors with missing data defined by the second bf() formula and X_2_ is a predictor or predictors without missing data. ‘Cormat’ represents the correlation matrix for the consensus phylogeny. Models were run for 8000 iterations. As a robustness check accounting for uncertainty in branch lengths, models were also run over a set of 100 phylogenies sampled from the posterior distribution of the phylogenetic inference analysis (described in detail in Supplementary Information) and the results combined using the combine_models() function in *brms*.

Sensitivity checks were run using additional measures of brain data and mantle length for species with measures from multiple individuals to ensure broadly similar results were attained. Sensitivity checks were also run with focal predictors with missing data in the main analyses (age at sexual maturity, predator breadth, diet breadth, hunting repertoire, and defense repertoire) with the missing cases dropped instead of imputed. These analyses did not contradict any of the findings from the imputed analyses (see analyses and supplementary data available at https://github.com/kcbasava/ceph-brain-evolution/tree/main/sensitivity-checks).

A detailed definition for each variable and operationalization in the analyses are provided in the Supplementary Information.

### Phylogeny

Although there are available phylogenies of coleoid cephalopod species (e.g. ^77,118^), the usual approach to comparative analyses would require matching the species in our dataset to those present on a chosen phylogeny and discarding those not included, which would significantly reduce our sample size. As we did not wish to discard any species with brain data, we decided to create an updated phylogeny for our analyses instead. With assistance from Jan Strugnell, we developed a new phylogeny for all 79 species by combining previously published phylogenetic analyses and additional sequence data on GenBank (described in Methods and in Basava et al.^119^).

In order to make use of all species for which brain measurements were available, we built a composite phylogenetic topology based on results of prior phylogenetic studies and inferred divergence times with the *BEAST2* software^120^ using sequence data from GenBank^121^. A detailed account of the methods for building the phylogeny are in the Supplementary Information. As relationships between coleoid species at more detailed levels than the octopodiforme-decapodiforme split are still uncertain^122,123^, we consider our phylogeny to be a rough approximation of a “true” set of evolutionary histories that nonetheless, when properly included in our statistical models, should guard against unobserved confounding through shared evolutionary trajectories. An ancestral state reconstruction of relative brain size (logged CNS/ML) using the ‘*contMap’* function from the package *phytools* in *R*^124^ is shown on the consensus phylogeny in Figure 6. Relative brain size is somewhat phylogenetically conserved, with larger measures appearing among the incirrate octopuses and cuttlefish.

## Author contributions

KB: Methodology, Software, Validation, Formal analysis, Investigation, Data curation, Writing – original draft, Writing – review & editing, Visualization, Project administration

TB: Conceptualization, Methodology, Software, Validation, Formal analysis, Investigation, Data curation, Writing – original draft, Writing – review & editing, Visualization, Supervision, Project administration

AL: Conceptualization, Methodology, Validation, Investigation, Data curation, Writing – review & editing, Visualization, Project administration

NG: Investigation, Data curation ZV: Investigation, Data curation JO: Investigation, Data curation

JM: Conceptualization, Methodology, Validation, Writing – review & editing, Supervision

MM: Conceptualization, Methodology, Validation, Software, Resources, Writing – original draft, Writing – review & editing, Supervision, Project administration, Funding acquisition

## Data Availability

Analysis code and data are openly available at www.github.com/kcbasava/ceph-brain-evolution. The pre-registration document can be found at www.osf.io/5dyfz.

## Acknowledgements

We thank the Templeton World Charity Foundation for supporting this work (TWCF0464). We would also like to thank Ryutaro Uchiyama for early feedback on the study design and Jan Strugnell for assistance and advice in compiling and constructing the phylogeny.

## Footnotes

1 For simplicity, we assume that brain size, complexity, and organization (e.g., neuronal density) are captured by a single state variable, which we will refer to as “size”.

2 Adaptive knowledge could potentially relate to locating food, avoiding predators, securing mates, hunting prey, and so on.

## Notes

https://cephdata.com

### Competing Interest Statement

The authors have declared no competing interest.

### Summary of Updates

Major revision to better emphasize the link between the data and underlying theory as well as clarifications for readability.

## References

1. Dunbar, R. I. M. The social brain hypothesis. Evol. Anthropol. 6, 178–190 (1998).

2. Rosati, A. G. Foraging Cognition: Reviving the Ecological Intelligence Hypothesis. Trends in Cognitive Sciences 21, 691–702 (2017).

3. Street, S. E., Navarrete, A. F., Reader, S. M. & Laland, K. N. Coevolution of cultural intelligence, extended life history, sociality, and brain size in primates. Proc. Natl. Acad. Sci. U.S.A. 114, 7908–7914 (2017).

4. Kaplan, H. S. & Robson, A. J. The emergence of humans: The coevolution of intelligence and longevity with intergenerational transfers. Proc. Natl. Acad. Sci. U.S.A. 99, 10221–10226 (2002).

5. Shou, W., Bergstrom, C. T., Chakraborty, A. K. & Skinner, F. K. Research: Theory, models and biology. eLife https://elifesciences.org/articles/07158 (2015) doi:10.7554/eLife.07158.

6. Servedio, M. R. et al. Not Just a Theory—The Utility of Mathematical Models in Evolutionary Biology. PLOS Biology 12, e1002017 (2014).

7. Muthukrishna, M., Doebeli, M., Chudek, M. & Henrich, J. The Cultural Brain Hypothesis: How culture drives brain expansion, sociality, and life history. PLOS Computational Biology 14, e1006504 (2018).

8. Fox, K. C. R., Muthukrishna, M. & Shultz, S. The social and cultural roots of whale and dolphin brains. Nature Ecology & Evolution 1, 1699–1705 (2017).

9. Aiello, L. C. & Wheeler, P. The Expensive-Tissue Hypothesis: The Brain and the Digestive System in Human and Primate Evolution. Current Anthropology 36, 199–221 (1995).

10. González-Forero, M., Faulwasser, T. & Lehmann, L. A model for brain life history evolution. PLoS Comput Biol 13, e1005380 (2017).

11. González-Forero, M. & Gardner, A. Inference of ecological and social drivers of human brain-size evolution. Nature 557, 554–557 (2018).

12. Isler, K. & Van Schaik, C. P. The Expensive Brain: A framework for explaining evolutionary changes in brain size. Journal of Human Evolution 57, 392–400 (2009).

13. Healy, S. D. & Rowe, C. A critique of comparative studies of brain size. Proc. R. Soc. B. 274, 453–464 (2007).

14. Lefebvre, L., Reader, S. M. & Sol, D. Brains, Innovations and Evolution in Birds and Primates. Brain Behav Evol 63, 233–246 (2004).

15. Reader, S. M. & Laland, K. N. Social intelligence, innovation, and enhanced brain size in primates. Proc. Natl. Acad. Sci. U.S.A. 99, 4436–4441 (2002).

16. Shultz, S. & Dunbar, R. I. M. Both social and ecological factors predict ungulate brain size. Proc. R. Soc. B. 273, 207–215 (2006).

17. Dunbar, R. I. M. & Shultz, S. Why are there so many explanations for primate brain evolution? Phil. Trans. R. Soc. B 372, 20160244 (2017).

18. Jolly, A. Lemur Social Behavior and Primate Intelligence: The step from prosimian to monkey intelligence probably took place in a social context. Science 153, 501–506 (1966).

19. Humphrey, N. K. The social function of intellect. in Growing points in ethology. (Cambridge U Press, Oxford, England, 1976).

20. Pérez-Barbería, F. J., Shultz, S. & Dunbar, R. I. M. EVIDENCE FOR COEVOLUTION OF SOCIALITY AND RELATIVE BRAIN SIZE IN THREE ORDERS OF MAMMALS. Evolution 61, 2811–2821 (2007).

21. Shultz, S. & Dunbar, R. I. M. Social bonds in birds are associated with brain size and contingent on the correlated evolution of life-history and increased parental investment: AVIAN BRAIN EVOLUTION. Biological Journal of the Linnean Society 100, 111–123 (2010).

22. Shultz, S. & Dunbar, R. I. M. Socioecological complexity in primate groups and its cognitive correlates. PHILOSOPHICAL TRANSACTIONS OF THE ROYAL SOCIETY B-BIOLOGICAL SCIENCES 377, (2022).

23. Machiavellian Intelligence: Social Expertise and the Evolution of Intellect in Monkeys, Apes, and Humans. (Clarendon Press ; Oxford University Press, Oxford : New York, 1988).

24. McNally, L., Brown, S. P. & Jackson, A. L. Cooperation and the evolution of intelligence. Proc. R. Soc. B. 279, 3027–3034 (2012).

25. Dávid-Barrett, T. & Dunbar, R. I. M. Processing power limits social group size: computational evidence for the cognitive costs of sociality. Proc. R. Soc. B. 280, 20131151 (2013).

26. Emery, N. J., Seed, A. M., Von Bayern, A. M. P. & Clayton, N. S. Cognitive adaptations of social bonding in birds. Phil. Trans. R. Soc. B 362, 489–505 (2007).

27. Iwaniuk, A. N. The Evolution of Cognitive Brains in Non-mammals. in Evolution of the Brain, Cognition, and Emotion in Vertebrates (eds Watanabe, S., Hofman, M. A. & Shimizu, T.) 101– 124 (Springer Japan, Tokyo, 2017). doi:10.1007/978-4-431-56559-8_5.

28. Hooper, R., Brett, B. & Thornton, A. Problems with using comparative analyses of avian brain size to test hypotheses of cognitive evolution. PLOS ONE 17, e0270771 (2022).

29. Pitnick, S., Jones, K. E. & Wilkinson, G. S. Mating system and brain size in bats. Proc. R. Soc. B. 273, 719–724 (2006).

30. Gonzalez-Voyer, A., Winberg, S. & Kolm, N. Social fishes and single mothers: brain evolution in African cichlids. Proc. R. Soc. B. 276, 161–167 (2009).

31. Finarelli, J. A. & Flynn, J. J. Brain-size evolution and sociality in Carnivora. Proc. Natl. Acad. Sci. U.S.A. 106, 9345–9349 (2009).

32. Kverková, K. et al. Sociality does not drive the evolution of large brains in eusocial African mole-rats. Sci Rep 8, 9203 (2018).

33. De Meester, G., Huyghe, K. & Van Damme, R. Brain size, ecology and sociality: a reptilian perspective. Biological Journal of the Linnean Society 126, 381–391 (2019).

34. Godfrey, R. K. & Gronenberg, W. Brain evolution in social insects: advocating for the comparative approach. J Comp Physiol A 205, 13–32 (2019).

35. O’Donnell, S. et al. Distributed cognition and social brains: reductions in mushroom body investment accompanied the origins of sociality in wasps (Hymenoptera: Vespidae). Proc. R. Soc. B. 282, 20150791 (2015).

36. Speechley, E. M., Ashton, B. J., Foo, Y. Z., Simmons, L. W. & Ridley, A. R. Meta-analyses reveal support for the Social Intelligence Hypothesis. Biological Reviews brv.13103 (2024) doi:10.1111/brv.13103.

37. Clutton-Brock, T. H. & Harvey, P. H. Primates, brains and ecology. Journal of Zoology 190, 309–323 (1980).

38. DeCasien, A. R., Williams, S. A. & Higham, J. P. Primate brain size is predicted by diet but not sociality. Nat Ecol Evol 1, 1–7 (2017).

39. Heldstab, S. A. et al. Manipulation complexity in primates coevolved with brain size and terrestriality. Sci Rep 6, 24528 (2016).

40. MacLean, E. L. et al. The evolution of self-control. Proceedings of the National Academy of Sciences 111, E2140–E2148 (2014).

41. Sol, D., Sayol, F., Ducatez, S. & Lefebvre, L. The life-history basis of behavioural innovations. Phil. Trans. R. Soc. B 371, 20150187 (2016).

42. González-Lagos, C., Sol, D. & Reader, S. M. Large-brained mammals live longer. J of Evolutionary Biology 23, 1064–1074 (2010).

43. Yu, X. et al. Large-brained frogs mature later and live longer: BRIEF COMMUNICATION. Evolution 72, 1174–1183 (2018).

44. Smeele, S. Q. et al. Coevolution of relative brain size and life expectancy in parrots. Proc. R. Soc. B. 289, 20212397 (2022).

45. Groot, N. E., Constantine, R., Garland, E. C. & Carroll, E. L. Phylogenetically controlled life history trait meta-analysis in cetaceans reveals unexpected negative brain size and longevity correlation. Evolution 77, 534–549 (2023).

46. Herrmann, E., Call, J., Hernández-Lloreda, M. V., Hare, B. & Tomasello, M. Humans have evolved specialized skills of social cognition:: The cultural intelligence hypothesis. SCIENCE 317, 1360–1366 (2007).

47. Whiten, A. & Van Schaik, C. P. The evolution of animal ‘cultures’ and social intelligence. Phil. Trans. R. Soc. B 362, 603–620 (2007).

48. Markov & Markov. Runaway brain-culture coevolution as a reason for larger brains: Exploring the “cultural drive” hypothesis by computer modeling. https://onlinelibrary.wiley.com/doi/full/10.1002/ece3.6350 (2020).

49. Doebeli, M., Hauert, C. & Killingback, T. The Evolutionary Origin of Cooperators and Defectors. Science 306, 859–862 (2004).

50. Wyles, J. S., Kunkel, J. G. & Wilson, A. C. Birds, behavior, and anatomical evolution. Proc. Natl. Acad. Sci. U.S.A. 80, 4394–4397 (1983).

51. Mather, J. A., Leite, T. S., Anderson, R. C. & Wood, J. B. Foraging and cognitive competence in octopuses. in Cephalopod Cognition (eds Darmaillacq, A.-S., Dickel, L. & Mather, J.) 125–149 (Cambridge University Press, 2014). doi:10.1017/CBO9781139058964.010.

52. Mather, J. A., Leite, T. S. & Batista, A. T. Individual prey choices of octopuses: Are they generalist or specialist? Current Zoology 58, 597–603 (2012).

53. Scheel, D. & Anderson, R. Variability in the diet specialization of Enteroctopus dofleini (Cephalopoda: Octopodidae) in the eastern Pacific examined from midden contents. American Malacological Bulletin 30, 267–279 (2012).

54. Brown, C., Garwood, M. P. & Williamson, J. E. It pays to cheat: tactical deception in a cephalopod social signalling system. Biol. Lett. 8, 729–732 (2012).

55. Hall, K. C. & Hanlon, R. T. Principal features of the mating system of a large spawning aggregation of the giant Australian cuttlefish Sepia apama (Mollusca : Cephalopoda). Marine Biology 140, 533–545 (2002).

56. Mather, J. Mating games squid play: reproductive behaviour and sexual skin displays in Caribbean reef squid Sepioteuthis sepioidea. Marine and Freshwater Behaviour and Physiology 49, 359–373 (2016).

57. Darmaillacq, A.-S., Dickel, L. & Mather, J. Cephalopod Cognition. (Cambridge University Press, 2014).

58. Mather, J. A. & Anderson, R. C. Exploration, play, and habituation in octopuses (Octopus dofleini). Journal of Comparative Psychology 113, 333–338 (1999).

59. Mather, J. A., Leite, T. S. & Batista, A. T. Individual prey choices of octopuses: Are they generalist or specialist? Current Zoology 58, 597–603 (2012).

60. Darmaillacq, A.-S., Dickel, L. & Mather, J. Cephalopod Cognition. (Cambridge University Press, 2014).

61. Hanlon, R. T. & Messenger, J. B. Cephalopod Behaviour. (2018).

62. Nixon, M. & Young, J. Z. The Brains and Lives of Cephalopods. (Oxford University Press, Oxford ; New York, 2003).

63. Mather, J. A. & Dickel, L. Cephalopod complex cognition. Current Opinion in Behavioral Sciences 16, 131–137 (2017).

64. Amodio, P. et al. Grow Smart and Die Young: Why Did Cephalopods Evolve Intelligence? Trends Ecol. Evol. 34, 45–56 (2019).

65. Schnell, A. K., Amodio, P., Boeckle, M. & Clayton, N. S. How intelligent is a cephalopod? Lessons from comparative cognition. Biol. Rev. 96, 162–178 (2020).

66. Ibáñez, C. M. & Keyl, F. Cannibalism in cephalopods. Rev Fish Biol Fisheries 20, 123–136 (2010).

67. Hanlon, R. T. & Messenger, J. B. Cephalopod Behaviour. (Cambridge University Press, 2018). doi:10.1017/9780511843600.

68. Mather, J. A. & Dickel, L. Cephalopod complex cognition. Current Opinion in Behavioral Sciences 16, 131–137 (2017).

69. Nixon, M. & Young, J. Z. The Brains and Lives of Cephalopods. (Oxford University Press, Oxford ; New York, 2003).

70. Amodio, P. et al. Grow Smart and Die Young: Why Did Cephalopods Evolve Intelligence? Trends in Ecology & Evolution 34, 45–56 (2019).

71. Mather, J. Where Should Comparative Cognition Be Going? To the Invertebrates. CCBR 19, 33–36 (2024).

72. Schnell, A. K., Amodio, P., Boeckle, M. & Clayton, N. S. How intelligent is a cephalopod? Lessons from comparative cognition. Biol Rev 96, 162–178 (2021).

73. Bendixen, T., Mather, J. & Muthukrishna, M. The Evolution of Big Brains: Toward a unifying theory. Inference: International Review of Science 6, (2021).

74. Packard, A. CEPHALOPODS AND FISH: THE LIMITS OF CONVERGENCE. Biological Reviews 47, 241–307 (1972).

75. Jaitly, R. et al. The evolution of predator avoidance in cephalopods: A case of brain over brawn? Front. Mar. Sci. 9, 909192 (2022).

76. Tanner, A. R. et al. Molecular clocks indicate turnover and diversification of modern coleoid cephalopods during the Mesozoic Marine Revolution. Proc. R. Soc. B. 284, 20162818 (2017).

77. Lindgren, A. R., Pankey, M. S., Hochberg, F. G. & Oakley, T. H. A multi-gene phylogeny of Cephalopoda supports convergent morphological evolution in association with multiple habitat shifts in the marine environment. BMC Evolutionary Biology 12, 129 (2012).

78. Woolley, S. N. C. et al. Deep-sea diversity patterns are shaped by energy availability. Nature 533, 393–396 (2016).

79. Angel, M. V. Biodiversity of the Pelagic Ocean. Conservation Biology 7, 760–72 (1993).

80. Danovaro, R., Snelgrove, P. V. R. & Tyler, P. Challenging the paradigms of deep-sea ecology. Trends in Ecology & Evolution 29, 465–475 (2014).

81. Costello, M. J. et al. A Census of Marine Biodiversity Knowledge, Resources, and Future Challenges. PLOS ONE 5, e12110 (2010).

82. Roper, C. F. E. & Voss, G. L. Guidelines for taxonomic descriptions of cephalopod species. he Biology and Resource Potential of Cephalopods: Memoirs of the National Museum of Victoria. http://repository.si.edu/xmlui/handle/10088/11335 (1983).

83. McClain, C. R. & Schlacher, T. A. On some hypotheses of diversity of animal life at great depths on the sea floor. Marine Ecology 36, 849–872 (2015).

84. Woolley, S. N. C. et al. Deep-sea diversity patterns are shaped by energy availability. Nature 533, 393–396 (2016).

85. Huffard, C. L. Ethogram of Abdopus aculeatus (d’Orbigny, 1834) (Cephalopoda: Octopodidae): can behavioural characters inform octopodid taxomony [taxonomy] and systematics? Journal of Molluscan Studies 73, 185–193 (2007).

86. Scata, G. Brain structure, body patterns and social interactions in the octopus Abdopus capricornicus. (The University of Queensland, 2022). doi:10.14264/053d1ea.

87. von Boletsky, S. & von Boletsky, M. V. Observations on the embryonic and early post-embryonic development of Rossia macrosoma (Mollusca, Cephalopoda). Helgolander Wissenschaftliche Meeresuntersuchungen 25, 135–161 (1973).

88. Montague, T. G., Rieth, I. J. & Axel, R. Embryonic development of the camouflaging dwarf cuttlefish, *SEPIA BANDENSIS* . Developmental Dynamics 250, 1688–1703 (2021).

89. Shultz, S. & Dunbar, R. I. M. Social bonds in birds are associated with brain size and contingent on the correlated evolution of life-history and increased parental investment: AVIAN BRAIN EVOLUTION. Biological Journal of the Linnean Society 100, 111–123 (2010).

90. Riverón, S. et al. Pelagic and benthic ecosystems drive differences in population and individual specializations in marine predators. Oecologia 196, 891–904 (2021).

91. Sayol, F. et al. Environmental variation and the evolution of large brains in birds. Nat Commun 7, 13971 (2016).

92. Ponte, G. et al. Cerebrotypes in Cephalopods: Brain Diversity and Its Correlation With Species Habits, Life History, and Physiological Adaptations. Front. Neuroanat. 14, 565109 (2021).

93. Navarrete, A. F., Reader, S. M., Street, S. E., Whalen, A. & Laland, K. N. The coevolution of innovation and technical intelligence in primates. Phil. Trans. R. Soc. B 371, 20150186 (2016).

94. Sol, D. Revisiting the cognitive buffer hypothesis for the evolution of large brains. Biol. Lett. 5, 130–133 (2009).

95. Kverková, K. et al. The evolution of brain neuron numbers in amniotes. Proc. Natl. Acad. Sci. U.S.A. 119, e2121624119 (2022).

96. Sol, D. et al. Neuron numbers link innovativeness with both absolute and relative brain size in birds. Nat Ecol Evol 6, 1381–1389 (2022).

97. Maddock, L. & Young, J. Z. Quantitative differences among the brains of cephalopods. Journal of Zoology 212, 739–767 (1987).

98. Wirz, K. Etude biometrique du Systeme nerveux des cephalopodes. Bull Biol 93, 78–117 (1959).

99. Smaldino, P. E. How to Translate a Verbal Theory Into a Formal Model. Social Psychology 51, 207–218 (2020).

100. Muthukrishna, M. & Henrich, J. A problem in theory. Nat Hum Behav 3, 221–229 (2019).

101. Borrelli, L. Testing the contribution of relative brain size and learning capabilities on the evolution of Octopus vulgaris and other cephalopods. (2007).

102. Dickel, L., Darmaillacq, A.-S., Jozet-Alves, C. & Bellanger, C. Learning, Memory, and Brain Plasticity in Cuttlefish (Sepia officinalis). in Handbook of Behavioral Neuroscience vol. 22 318–333 (Elsevier, 2013).

103. Liu, Y. et al. Morphological changes of the optic lobe from late embryonic to adult stages in oval squids *Sepioteuthis lessoniana*. Journal of Morphology 279, 75–85 (2018).

104. Ovalle, F. V., Navarrete, A. G. & Castillo, H. S. Characterization of the brain of the Red Mayan octopus (Octopus Maya). Preprint at 10.21203/rs.3.rs-1805491/v1 (2022).

105. Chung, W.-S., Kurniawan, N. D. & Marshall, N. J. Comparative brain structure and visual processing in octopus from different habitats. Current Biology 32, 97–110.e4 (2022).

106. Chung, W.-S., López-Galán, A., Kurniawan, N. D. & Marshall, N. J. The brain structure and the neural network features of the diurnal cuttlefish Sepia plangon. iScience 26, 105846 (2023).

107. Montague, T. G. et al. A brain atlas for the camouflaging dwarf cuttlefish, Sepia bandensis. Current Biology 33, 2794–2801.e3 (2023).

108. Cephalopod Life Cycles. (Academic Press, London ; New York, 1983).

109. Boyle, P. R. & Rodhouse, P. Cephalopods ecology and fisheries. in Cephalopods. Ecology and Fisheries (Blackwell Science, Ames, Iowa, 2005). doi:10.1002/9780470995310.

110. Jereb, Roper, Norman & Finn. Cephalopods of the World Vol. 3. (FOOD & AGRICULTURE ORG, Place of publication not identified, 2014).

111. Cephalopods of the World Vol. 1. (Food and Agriculture Organization of the United Nations, Rome, 2005).

112. Bürkner, P.-C. brms: An R Package for Bayesian Multilevel Models Using Stan. J. Stat. Soft. 80, (2017).

113. Carpenter, B., et al. *Stan* : A Probabilistic Programming Language. J. Stat. Soft. 76, (2017).

114. McElreath, R. *Statistical Rethinking: A Bayesian Course with Examples in R and Stan*. (Taylor and Francis, CRC Press, Boca Raton, 2020).

115. Kurz, A. S. Statistical Rethinking with Brms, Ggplot2, and the Tidyverse: Version 1.0.1. (2019).

116. Greenland, S., Pearl, J. & Robins, J. M. Causal Diagrams for Epidemiologic Research. Epidemiology 10, 37–48 (1999).

117. Bürkner, P. Estimating Phylogenetic Multilevel Models with brms. brms: Bayesian regression models using Stan https://paul-buerkner.github.io/brms/articles/brms_phylogenetics.html#a-simple-phylogenetic-model (2024).

118. López-Córdova, D. A. et al. Mesozoic origin of coleoid cephalopods and their abrupt shifts of diversification patterns. Molecular Phylogenetics and Evolution 166, 107331 (2022).

119. Basava, K. et al. A phylogeny of extant coleoid cephalopods with brain data. Preprint at 10.1101/2024.04.29.591691 (2024).

120. Bouckaert, R. et al. BEAST 2.5: An advanced software platform for Bayesian evolutionary analysis. PLoS Comput Biol 15, e1006650 (2019).

121. Benson, D. A. et al. GenBank. Nucleic Acids Research 41, D36–D42 (2012).

122. Lindgren, A. R., Pratt, A., Vecchione, M. & Anderson, F. E. Finding a home for the ram’s horn squid: phylogenomic analyses support Spirula spirula (Cephalopoda: Decapodiformes) as a close relative of Oegopsida. Org Divers Evol 23, 91–101 (2023).

123. Sanchez, G. et al. Genus-level phylogeny of cephalopods using molecular markers: current status and problematic areas. PeerJ 6, e4331 (2018).

124. Revell, L. J. phytools: an R package for phylogenetic comparative biology (and other things). Methods in Ecology and Evolution 3, 217–223 (2012).

